# Diverse Visual Experience Promotes Integrated and Human-Aligned Face Representations in Deep Neural Networks

**DOI:** 10.64898/2025.12.19.695398

**Authors:** Elaheh Akbari, Katharina Dobs

## Abstract

Humans are experts at recognizing faces, yet this expertise is not uniform: people perceive faces from familiar facial groups more accurately than those from unfamiliar ones—a phenomenon known as the Other-Race Effect (ORE). Diverse facial exposure mitigates this bias, but how it reorganizes face representations to support cross-group recognition remains unclear. To address this question, we combined controlled training variation in deep convolutional neural networks (CNNs) with analyses of representational geometry, lesioning, and human behavioral data. CNNs trained exclusively on either Asian or White faces reproduced ORE-like biases, whereas a CNN trained on both groups showed reduced bias and balanced recognition performance. Critically, representational similarity analyses and lesioning revealed largely overlapping feature spaces across groups, indicating an integrated representational organization. Trial-by-trial comparisons with human choices further showed that the dual-trained CNN best captured human face-matching behavior across both Asian and White participants, outperforming single-trained networks on cross-group trials. Together, these findings show that diverse visual experience promotes an integrated representational geometry that supports cross-group generalization, providing a computational account of how exposure diversity may shape human face representations.

## Introduction

Humans are remarkably proficient at visual recognition, yet our perception consistently reflects systematic biases. These biases shape how we interpret sensory information, ranging from low-level visual illusions, such as perceiving motion in static images or misjudging size and color based on context, to higher-level biases like better recognition of one’s own social group. One prominent example is the Other-Race Effect (ORE), the difficulty humans experience when recognizing faces from populations less familiar to them. The ORE is robust and extensively documented across cultures and contexts (1–3), in experimental paradigms such as memory and perceptual discrimination (4–8), as well as in neuroimaging studies (9, 10).

Developmental research suggests that the ORE emerges from asymmetrical visual experience: although absent at birth (11), own-group preferences arise by 3 months and persist into adulthood (12–14). Early exposure to diverse faces can significantly mitigate the ORE (12, 15–20), a pattern supported by behavioral (21, 22), computational (23), and neuroimaging evidence (24). Importantly, recent findings show that not only which faces one encounters, but also the richness of face experience shapes recognition abilities: individuals from smaller or less socially diverse communities show reduced face recognition performance (25). Together, these results suggest that the diversity of one’s visual environment broadly constrains the development of robust face representations. Despite these findings, how visual experience affects the underlying representational geometry of faces remains unclear. In particular, does limited experience compress the representational space needed to individuate unfamiliar faces, and does diverse exposure promote segregated group-specific representations or a more integrated representational space that supports cross-group generalization? Addressing these questions in humans is challenging due to limited control over individual perceptual histories and the difficulty of disentangling perceptual from social factors.

Artificial neural networks—lacking social cognition and motivation—provide a powerful framework for isolating the role of visual experience in shaping face perception. Deep convolutional neural networks (CNNs), in particular, have proven valuable models of biological vision: when trained on object recognition, they exhibit strong similarities to the primate visual system in both performance and representational geometry (26–28). Moreover, CNNs reproduce several hallmark signatures of human face perception, including the inversion effect (29, 30), Mooney face perception (31), and the ORE itself (29, 32). Recent work further shows that CNNs trained jointly on face and object recognition spontaneously develop segregated subsystems for each task (33), mirroring functional specialization for faces and objects in the human visual cortex. Lesioning analyses reveal a double dissociation: selectively ablating the face-processing subnetwork impairs face recognition but not object recognition, and vice versa, indicating segregated domain-specific feature spaces. Manipulating visual experience in CNNs also shows that exposure to faces is necessary for view-invariant representations (34), and that both the amount and type of facial experience shape view invariance and sensitivity to diagnostic face features (35). Together, these findings highlight key methodological advantages of CNNs: their training ‘diet’ can be precisely controlled, and their internal representations can be causally probed via lesioning, enabling systematic tests of how visual experience shapes representational geometry (36–39).

Here, we leverage CNNs to investigate how exposure diversity shapes face representations and contributes to the ORE. Specifically, we asked whether training diversity promotes segregated group-specific representations or a more integrated representational space, and whether the resulting latent representations predict human face perception behavior across groups. To address these questions, we trained three CNNs: one only on Asian faces, one only on White faces, and one on both groups. We first replicated that single-trained CNNs show an ORE and that diverse training reduces this bias. We then examined how these differences are reflected in the networks’ internal representational geometry and how well each model accounts for human behavioral data. We found that single-trained CNNs developed group-specific feature spaces that were compressed for the unfamiliar group. In contrast, diverse training yielded more integrated representations that aligned with human behavior across groups. By combining computational modeling with human behavioral data, our work provides a computational account of how visual experience shapes face representations and contributes to perceptual bias, offering insights into how diverse experience may support more generalized face perception in humans.

## Results

### ORE-like biases emerge in single-trained CNNs and are reduced by diverse training

To investigate how training diversity influences recognition biases, we trained three CNNs based on VGG16 architecture (40): Single White (trained on only White faces), Single Asian (trained on only Asian faces), and Dual CNN (trained on both Asian and White faces). We evaluated the models on an identity-matching task using a novel set of 40 White and 40 Asian facial identities (five images per identity; Figure 4C) that were not included in training. We verified that the two image sets did not differ in global pixel variance (White: mean = .073, Asian: mean = .073; p = .84; bootstrap test), reducing the likelihood that low-level image statistics contributed to performance differences. Using this test set, we confirmed the presence of an ORE in the single-trained CNNs. Both networks performed significantly better on faces from their trained group than on faces from the untrained group, as measured by face-matching accuracy in the target-matching task (Single Asian CNN: Asian-face accuracy = .94 vs. White-face accuracy = .90; Single White CNN: Asian-face accuracy = .90 vs. White-face accuracy = .97; all *p* < .0001, Bonferroni-corrected; Figure 1A), confirming a clear ORE. In line with previous findings (29, 32), these results demonstrate that asymmetric visual experience alone is sufficient to produce group-specific recognition biases.

**Figure 1.**
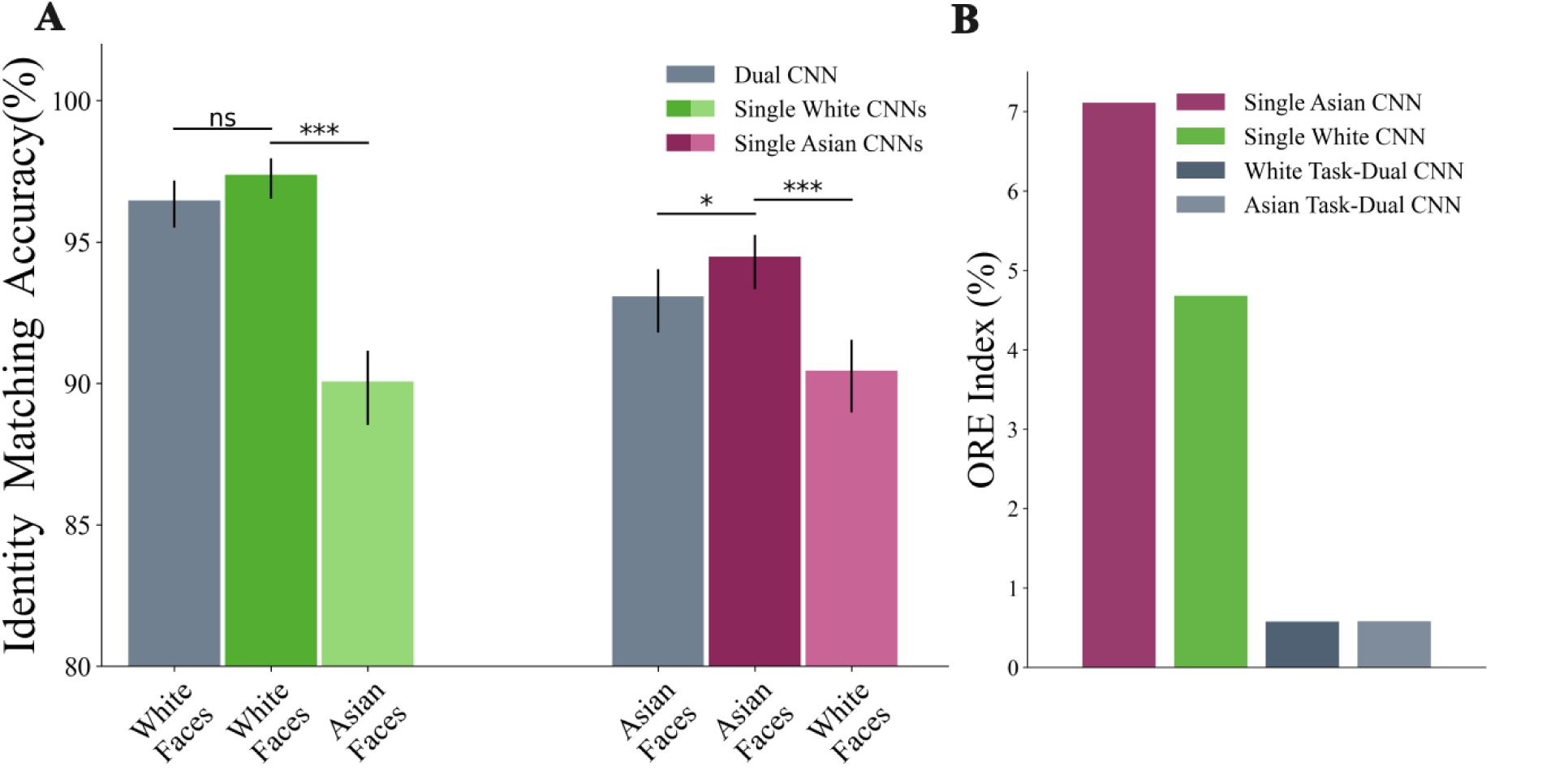
Recognition biases in CNNs. A) Identity-matching accuracy for White and Asian faces across three CNNs: Single White (green), Single Asian (purple), and Dual (gray) CNN. All pairwise comparisons were statistically significant (p < .05, Bonferroni-corrected), except for the comparison between the Dual CNN and the Single White CNN on White faces (p = .06, Bonferroni-corrected). Error bars represent 95% bootstrapped confidence intervals. B) ORE index for each model. Single-trained CNNs exhibit substantial bias favoring their trained facial group, whereas the Dual CNN shows minimal ORE, indicating balanced recognition performance across facial groups. Asterisks indicate significance level as follows: ns, not significant; *, p < .05; ***, p < .0001.

Next, we confirmed that training diversity mitigates these biases while preserving high performance across both facial groups. The Dual CNN, trained on both facial groups, maintained high accuracy for White and Asian faces (Asian = .93, White = .96, *p* < .0001; Figure 1A). Notably, its overall accuracy across groups (mean = .95) exceeded that of either single-trained model (Single Asian: mean = .92; Single White: mean = .94), indicating that diverse experience not only balances performance but also improves it. Its performance on White faces did not differ significantly from that of the Single White CNN (*p* = .06), whereas the Single Asian CNN slightly but significantly outperformed the Dual CNN on Asian faces (*p* = .03). Crucially, this balanced performance is particularly impressive given that the Dual CNN was trained on only half as many images per identity as the single-trained CNNs, indicating that training diversity, rather than data quantity, drives its broader generalization.

To further quantify these recognition biases, we computed an ORE index for each CNN (Figure 1B), measuring the relative drop in recognition accuracy for untrained versus trained groups (higher values correspond to stronger bias). The results mirrored the accuracy pattern observed above: single-trained CNNs showed larger ORE indices, indicating greater difficulty recognizing untrained facial groups (Single White CNN = 4.7%; Single Asian CNN = 7.1%). In contrast, the Dual CNN demonstrated minimal bias, with a baseline-weighted ORE index of ∼0.6% for both groups. These results underscore the effectiveness of diverse training in mitigating representational bias. Importantly, this pattern parallels developmental findings in humans, where early exposure to diverse faces has been shown to reduce the ORE (12, 15–19, 21, 22).

### Lesioning reveals an integrated feature space in the Dual CNN

The performance results showed that training diversity substantially reduced recognition biases in the Dual CNN. We next asked how this bias reduction is reflected in the networks’ internal representations. Specifically, we assessed whether the Dual CNN’s reduced ORE arises from a shared, integrated representational space that encodes faces across groups, or from the coexistence of separate group-specific subspaces supporting each facial group, as previously found for faces and objects (33). To test this, we performed a lesioning analysis targeting the most critical filters in the Dual CNN (Figure 2A). In CNNs, filters are spatially localized feature detections—sets of weights that respond to specific visual patterns in the input image. Filters were ranked by the increase in loss caused by ablating them during identity recognition for either facial group. We then lesioned the top 20% of filters most critical for each group-specific task (e.g., recognition of all Asian identities) and measured the resulting change in validation performance.

**Figure 2.**
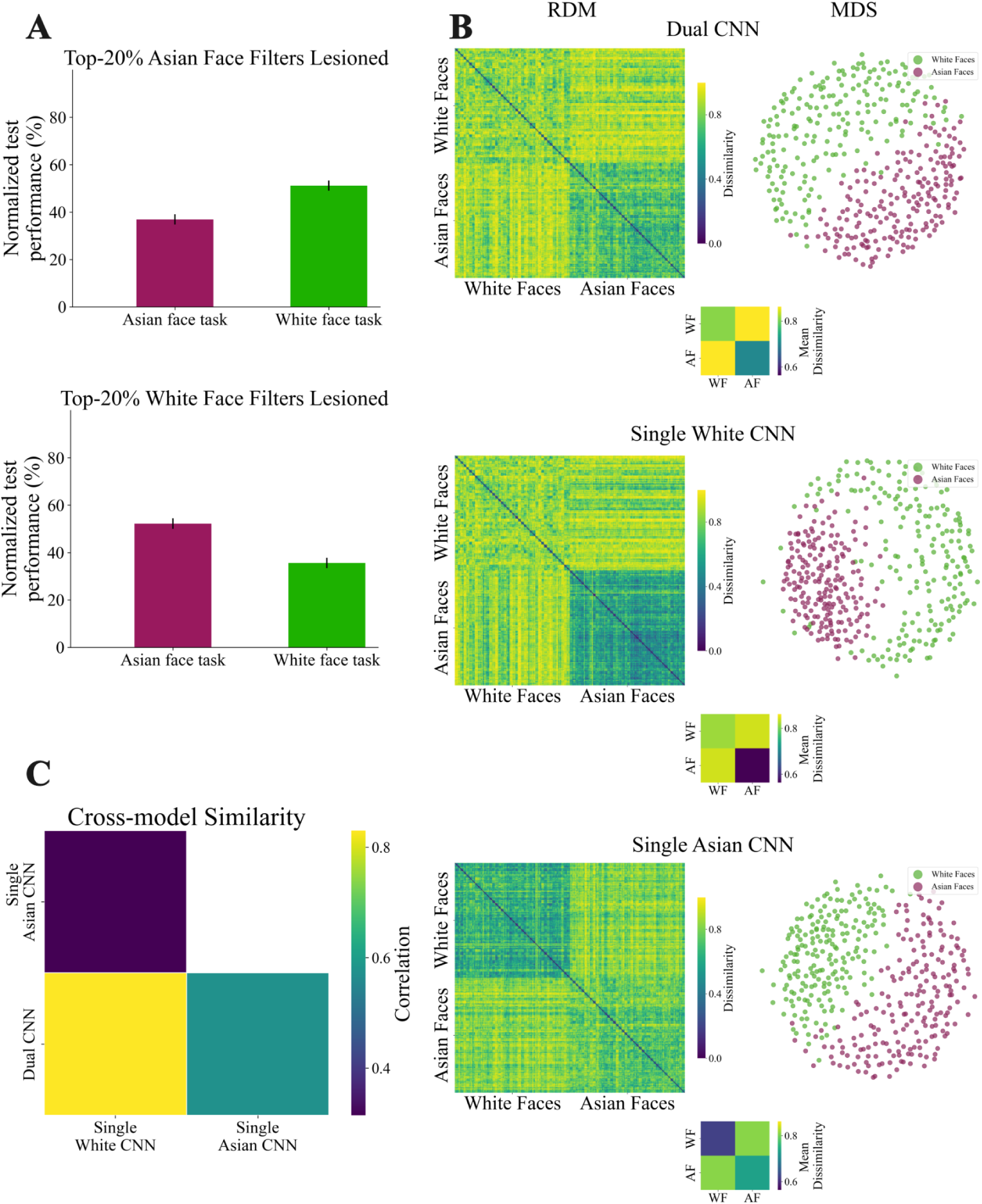
Properties of the representational feature spaces. A) Lesioning analysis. The top 20% of group-specific filters were ablated, and performance was evaluated across both facial groups. Lesioning impaired not only the target group but also the non-target group, indicating shared representational resources within the Dual CNN. Performance is shown relative to each task’s baseline accuracy. Error bars represent 95% bootstrapped confidence intervals. B) Representational Dissimilarity Matrices (RDMs). Pairwise dissimilarities (1 – cosine similarity) among White and Asian face identities for each CNN. To the right of each RDM, we show (i) MDS embedding of the dissimilarities (White faces: green; Asian faces: purple) and (ii) a 2×2 mean-dissimilarity plot quantifying within– and between-group similarity (AF: Asian faces; WF: White faces). Single-trained CNNs exhibit lower within-group dissimilarity for the untrained group, indicating reduced identity discriminability. In contrast, the Dual CNN shows balanced dissimilarity patterns across both groups, consistent with a more integrated representational space. C) Cross-model RSA. Using Pearson correlation to compare representational geometries across CNNs.

Lesioning group-specific filters in the Dual CNN resulted in substantial declines in performance across both tasks (lesioning White-selected filters: White faces: –65%, Asian faces: –48%; Lesioning Asian-selected filters: White faces: –49%, Asian faces: –63%). Notably, 64% of filters were shared between the two sets of top-ranked filters (65 out of 102 filters per task), indicating considerable overlap in the filters most critical for each facial group. Although impairments were somewhat larger for the facial group corresponding to the lesioned filters, both lesion types strongly disrupted performance for both groups. Together, these results indicate that the Dual CNN relies on largely shared representational resources for identity recognition across facial groups, consistent with an integrated rather than segregated representational organization.

### Training diversity leads to balanced representational geometry

To further probe the structure of this shared representational space and compare it with the single-trained CNNs, we examined the geometry of their feature spaces using Representational Similarity Analysis (RSA). For each model, we first constructed representational dissimilarity matrices (RDMs) by computing pairwise cosine similarities between activation patterns for all face identities within and between facial groups (Figure 2B; RDMs). In the Single White CNN, within-Asian similarity was significantly higher than within-White similarity (mean = .43 vs. .18; *p* < .0001; bootstrap test), indicating lower discriminability among Asian identities—consistent with the previously observed ORE. Similarly, in the Single Asian CNN, within-White similarity exceeded within-Asian similarity (mean = .37 vs. .26; *p* < .0001), aligning with reduced recognition of White faces (Figure 2B; mean-dissimilarity plots). This is supported by Multidimensional Scaling (MDS) visualizations, where the trained groups are dispersed but the untrained groups are compressed in each single-trained CNN, indicating poor discriminability for the untrained group compared to the trained group. On the other hand, in the Dual CNN, the similarities are evenly distributed, reflecting shared representational space with comparable discriminability for both groups (Figure 2B; MDS plots).

Interestingly, the Dual CNN displayed within-group similarity comparable to each of the corresponding single-trained models (within-Asian similarity: Dual = .29 vs. Single Asian = .26; *p* < .001; within-White similarity: Dual = .19, Single White CNN = .18, *p* < .05), reflecting a more uniform representational structure characterized by evenly distributed identity clusters. Analyses of between-group similarity revealed that the Dual CNN organized the two facial groups into more distinct but coexisting subspaces (between-group similarity = .13, compared to the Single Asian CNN = .19 and the Single White CNN = .15; all pairwise *p*s < .0001; Bonferroni-corrected) (Figure 2B; mean-dissimilarity plots). These findings suggest a hierarchically integrated representational organization: the Dual CNN encodes both groups within a common feature space that supports cross-group generalization, yet preserves sufficient separation to maintain high identity performance (see Figure 1A) for each facial group. Consistent with this interpretation, cross-model RSA showed that the Dual CNN’s representational geometry was more similar to both single-trained networks (Single White CNN: *r* = .83; Single Asian CNN: *r* = .57; all pairwise *p*s < .0001; bootstrap test, Bonferroni-corrected) than the single-trained models were to each other (*r* = .31, *p* < .0001), with a somewhat stronger resemblance to the White-trained network (Figure 2C). This pattern further supports the view that the Dual CNN developed an integrated, but structured, representational space that bridges both facial domains.

Together, these results highlight a shared conclusion: training diversity promotes the emergence of integrated and generalizable feature representations. These representations are less constrained by group-specific biases, supporting more balanced identity encoding across facial groups and enabling improved cross-group face recognition performance in the Dual CNN.

### Dual CNN shows the most consistent correspondence with human behavior across conditions

To evaluate the behavioral relevance of these representational differences, we next compared the CNNs’ decision patterns to human face-matching behavior. We used previously collected behavioral data from Asian (*n* = 102) and White (*n* = 269) participants performing a target-matching task (29). Consistent with prior work, humans showed a modest but significant ORE, with performance decreasing by 4.2% for White participants and 3.0% for Asian participants on other-group triplets. To compare human and model behavior, we computed Pearson correlations between each CNN’s predictions and the average human responses for each stimulus triplet in the target-matching task (Figure 4D). This analysis quantified how well each CNN captured human choice patterns across stimuli (Figure 3). A noise ceiling was estimated using split-half reliability across participants (Spearman–Brown corrected) to determine the upper bound of explainable variance.

**Figure 3.**
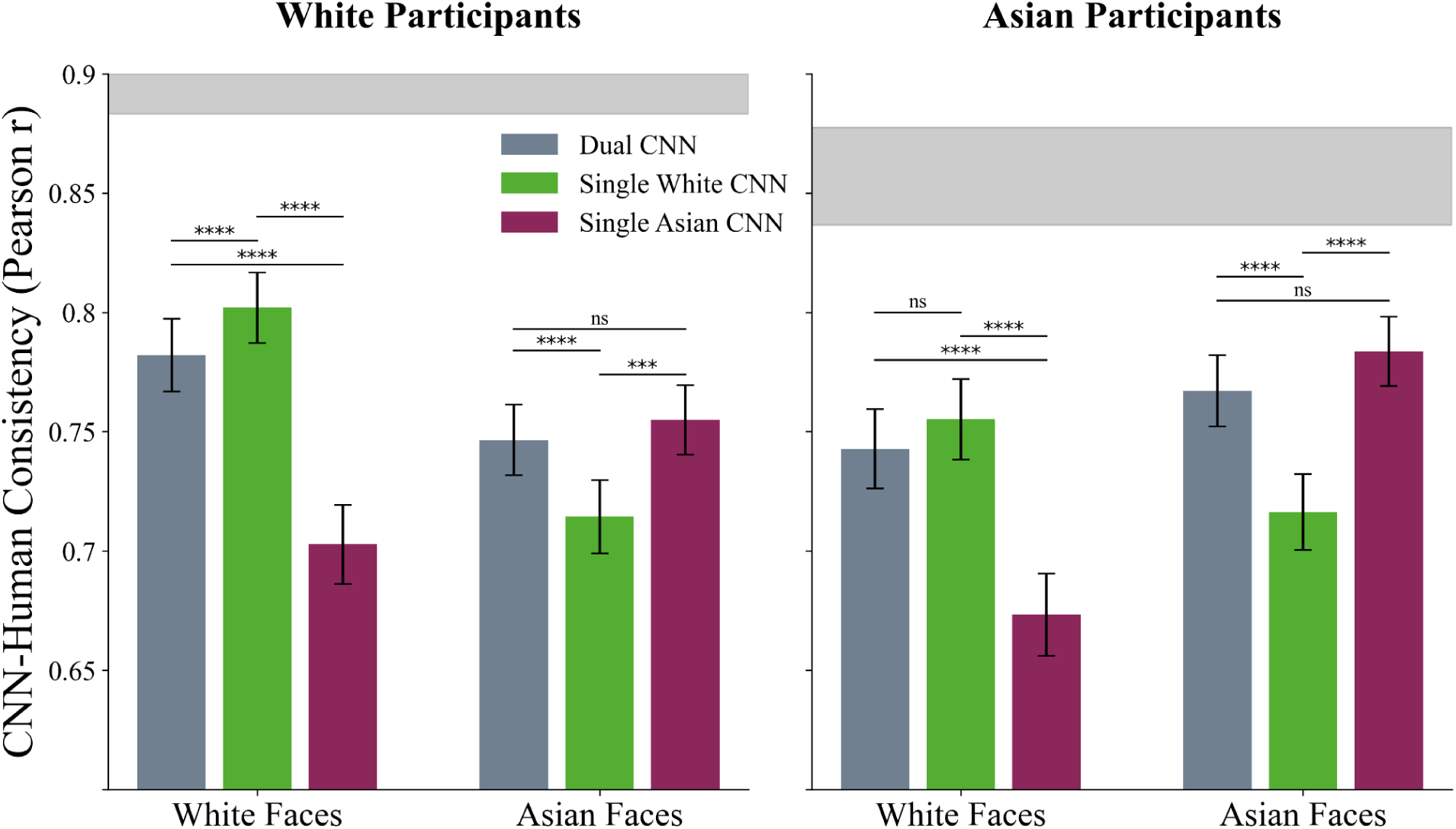
Behavioral consistency between human participants and CNNs. CNNs’ choice probabilities (see Methods) were correlated with the mean human responses across participants, computed separately for White (left panel; n = 269) and Asian (right panel; n = 102) participant groups. Error bars represent bootstrapped standard deviations of the correlation coefficients. Gray horizontal bars indicate Spearman-Brown-corrected noise ceilings. Asterisks denote significance levels: ns, not significant (p > 0.05); ** p < .001; *** p < .0001.

**Figure 4.**
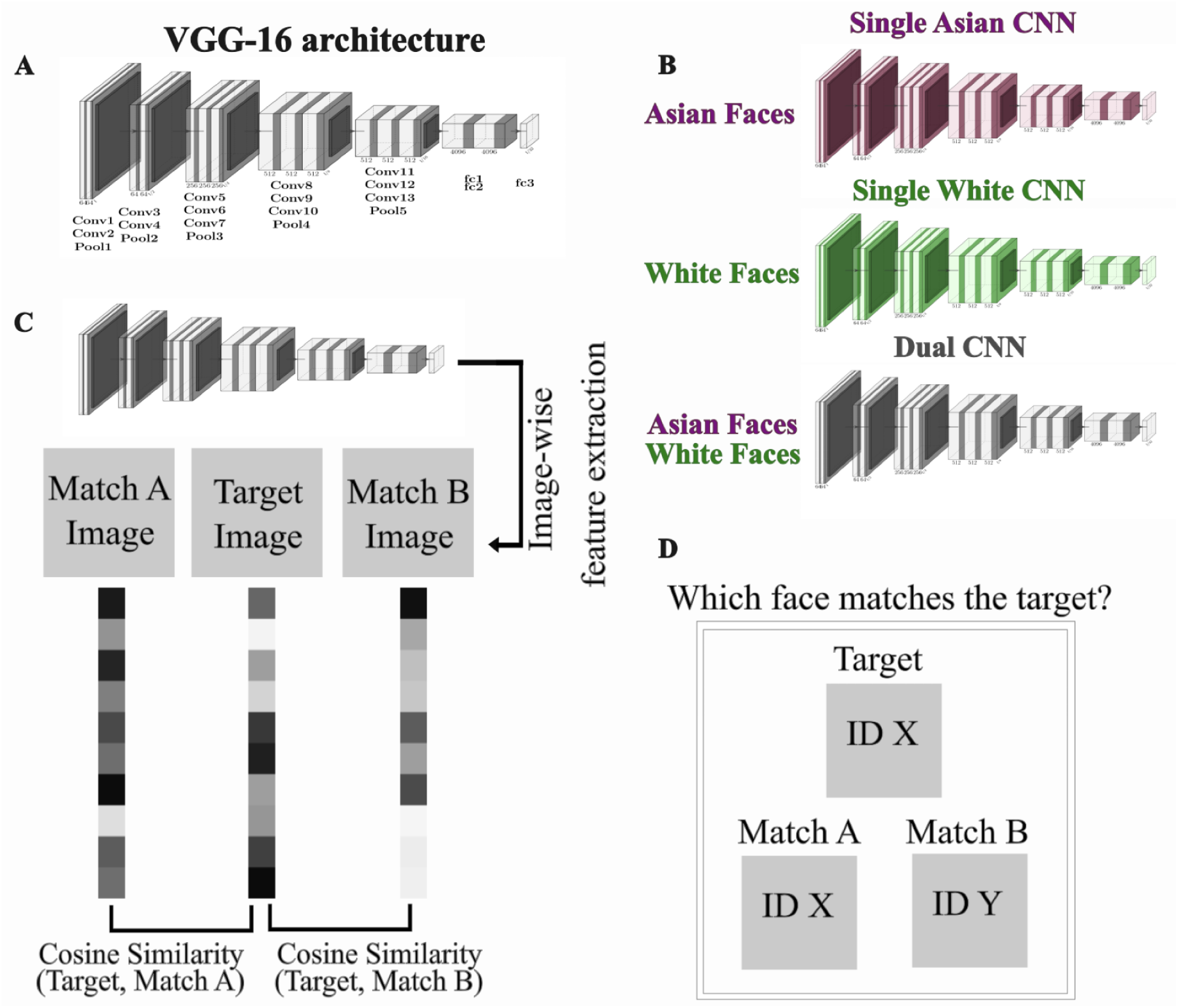
Overview of the methods. A) VGG16 architecture consisting of 13 convolutional, 5 pooling, and 3 fully connected layers. B) Overview of trained CNNs: a Single Asian CNN trained only on Asian identities, a Single White CNN trained only on White identities, and a Dual CNN trained on both Asian and White identities. C) Target-matching task used in CNNs. Image-wise feature vectors were extracted from the penultimate fully connected layer of each CNN, and cosine similarity was computed between the target image and each match (Match A and Match B). This cosine similarity is subsequently used to compute accuracy and choice probability (see Methods for details). D) Human target-matching task, in which participants chose whether Match A or Match B depicted the same identity as the target face. Images displayed here are not part of the stimulus set. Images are publicly available at https://commons.wikimedia.org.

For White participants, the Single White CNN best predicted behavioral responses on White-face trials (*r* = .80, *p* < .0001; bootstrap test), followed by the Dual CNN (*r* = .78, *p* < 0.0001) and the Single Asian CNN (*r* = .70, *p* < 0.0001). The Single White CNN correlated significantly more strongly with human behavior than either the Dual CNN or the Single Asian CNN (all *p* < .0001; Bonferroni-corrected). When White participants viewed Asian faces, behavior was best explained by both the Single Asian CNN (*r* = .75, *p* < .0001) and the Dual CNN (*r* = .75, *p* < .0001), with no significant difference between the two (*p* = 1). Both, however, predicted behavior significantly better than the Single White CNN (*r* = .71; *p* < .0001; *p* < .001 vs. Single Asian; *p* < .0001 vs. Dual).

For Asian participants, a similar group-specific pattern emerged. On Asian-face triplets, both the Single Asian CNN (*r* = .78, *p* < .0001; bootstrap test) and the Dual CNN (*r* = .77, *p* < .0001) showed strong correspondence with human behavior and did not differ significantly from each other (*p* = .056; Bonferroni-corrected). In contrast, when Asian participants viewed White faces, the Single White CNN (*r* = .75, *p* < .0001) and the Dual CNN (*r* = .74, *p* < .0001) provided a better fit than the Single Asian CNN (*r* = .67 ; *p* < .0001). The Single White CNN and Dual CNN performed comparably (*p* = .1) and both outperformed the Single Asian CNN (both *p* < .0001). Across both participant groups, this asymmetric generalization pattern likely reflects differences in real-world visual experience: unlike the single-trained CNNs, which never encountered the other facial group, human observers have at least some diverse exposure (25). These differences help explain why Asian participants’ cross-group choices align better with the Single White CNN than with the Single Asian CNN, and why White participants’ cross-group choices align better with the Single Asian CNN than with the Single White CNN: in each case, the model trained on a given facial group better approximates observers’ greater relative experience with that group than a model with no exposure to it.

Taken together, the single-trained CNNs aligned best with participants who were most familiar with the facial group each model was trained on, showing the strongest correspondence on own-group faces and capturing a substantial portion of the explainable variance in human behavior. Importantly, the Dual CNN performed comparably to the single-trained CNNs across all conditions, even outperforming them on cross-group trials. This pattern indicates that diverse training fosters more flexible, generalizable representations, yielding a model whose decision patterns more closely reflect human face processing across groups.

## Discussion

Our aim in this study was to investigate how diversity in visual experience shapes the representational geometry of faces and supports cross-group recognition. To address this question, we used CNNs as fully controlled and accessible models of visual experience. Consistent with prior findings, CNNs trained exclusively on a single face set exhibited a clear ORE, whereas the Dual CNN showed markedly reduced bias and robust identity recognition across both groups. Lesioning and representational similarity analyses revealed that this mitigation does not arise from segregated group-specific subsystems but from an integrated representational space that relies on largely overlapping feature sets to individuate faces across groups. Critically, the Dual CNN also showed the strongest correspondence with human face-matching behavior across participant groups, suggesting that diverse visual experience promotes representational structures that more closely align with human perceptual strategies.

Our findings are consistent with developmental studies in humans showing that early exposure to diverse faces is associated with reduced ORE and individuated processing of less familiar groups (12, 15–19, 21, 22). Similarly, the Dual CNN developed more generalizable face representations, suggesting that diversity during training, whether biological or artificial, plays a critical role in shaping the flexibility of face-identity processing systems. Importantly, in humans, the timing of exposure appears crucial: diverse experience early in development has a stronger mitigating effect than exposure later in life (17). Future work could investigate whether artificial systems exhibit analogous “developmental windows”.

Previous computational studies have documented ORE-like biases in CNNs trained on homogeneous face datasets (29, 32, 41). Building on this work, our results replicate these effects but extend them in several key ways. First, we systematically compared models trained on fully controlled training regimes, isolating the contribution of training diversity with fewer confounds than in earlier work. Second, by integrating lesioning with representational similarity analyses, we moved beyond correlational descriptions of representational geometry to causally test whether the features supporting recognition are shared or group-specific, allowing us to distinguish integrated from segregated representational geometries. Third, by comparing model predictions with human behavioral data across two participant groups, we show that the Dual CNN provides the strongest overall correspondence with human performance, underscoring the relevance of these computational mechanisms for understanding human perception.

The lesioning results address a broader mechanistic question regarding the conditions under which functional segregation emerges. We found that the majority of the critical filters were shared across facial groups, and that lesioning group-selected filters impaired performance for both tasks. This pattern indicates that optimization over subgroups within a single domain (White and Asian faces) does not promote strongly segregated feature spaces. In contrast, prior work has shown that optimization over distinct domains (e.g., faces vs. objects) results in functionally segregated subsystems (33, 42, 43). Together, these findings suggest that functional segregation may arise primarily when computational demands require qualitatively different feature sets, whereas optimization across subgroups of a single domain promotes the reuse and flexible weighting of shared features.

The representational geometry observed across models also aligns with the “face space” theory (44), which posits that other-race faces are represented in a more compact and thus less informative representational space. Recent EEG and neural decoding studies support this account, showing that other-race faces elicit less discriminable neural signals and homogeneous representations (45, 46). This mirrors the compressed and less separable representation observed in the single-trained CNNs. In contrast, the Dual CNN showed that training diversity expands the “face space” enabling finer-grained and more structured identity representations across groups. This broadened representational geometry is consistent with recent computational work showing that richer and more varied visual training improves the separability and organization of learned object representations (47, 48).

Together, these results support an interpretation of the ORE as a form of representational overfitting when exposure is restricted to a narrow subset of faces: the system allocates representational resources to maximize discrimination within the familiar group, at the cost of compressed and less informative representations of unfamiliar groups. Importantly, such specialization may not be purely detrimental. In biological systems, this tuning may reflect an adaptive optimization to the statistics of one’s social environment, enhancing discrimination for the identities observers most frequently need to individuate. In this sense, the ORE may constitute a “feature” of efficient resource allocation, albeit one that becomes a “bug” in modern multicultural environments where cross-group recognition is increasingly important. Future work should investigate the benefits and costs associated with maintaining a shared versus specialized feature space, and may draw parallels to other domains such as language, where bilingualism also shows complex trade-offs (49).

Human-model comparisons also revealed a meaningful asymmetry between artificial training and modern human experience. Single-trained CNNs best captured participant choices for their own-group faces, but on other-group trials, human behavior aligned more closely with the Dual CNN or with the CNN trained on the corresponding group. Unlike CNNs exposed exclusively to one facial group, human observers, no matter how limited their reported contact, inevitably accumulate heterogeneous visual input from indirect sources such as media, advertising, or incidental encounters. Such incidental exposure can contribute to recognition ability (25), whereas self-reported contact explains only modest variance in other-group performance (50). More broadly, our findings indicate that cross-group performance reflects not only the observer’s perceptual history, but also the structure of the recognition task itself. Depending on which visual cues are diagnostic in a given trial, task demands can either accentuate or attenuate group differences in both humans and models.

Our work also provides insights into the potential origins of the ORE by showing that perceptual-expertise mechanisms (51) are sufficient to produce ORE-like biases in a system optimized solely for face recognition. Because CNNs lack any socio-motivational constructs, the emergence of ORE in these models must arise from perceptual learning dynamics alone, specifically from asymmetrical experience. Moreover, we show that this bias can be substantially mitigated when the model receives balanced, diverse facial input. Together, these findings suggest that the human ORE can, in principle, be explained by perceptual factors alone, while socio-cognitive factors (52, 53) may modulate or amplify this bias rather than generate it de novo. This challenges pure socio-cognitive explanations and instead supports hybrid accounts in which perceptual-expertise mechanisms provide the foundational structure on which socio-cognitive influences act (9, 54).

From a machine learning standpoint, our findings echo broad concerns in the AI fairness literature: models trained on imbalanced datasets not only underperform on unfamiliar inputs, but also exhibit amplified representational biases. This is consistent with prior work showing how skewed training distributions can exacerbate bias in computer vision systems (32, 55, 56). Such imbalances can lead to poorly generalizable models that behave unreliably or unfairly when deployed in real-world settings. By showing that a model trained on a more diverse dataset (Dual CNN) improves performance across groups and develops more robust, generalizable representational geometry, our results reinforce calls in the AI community for proactive dataset diversification as a critical step toward building fairer and more reliable vision systems.

This study has several limitations. First, all analyses were performed using a feedforward CNN architecture. Although well established for face recognition (57), it remains to be tested whether similar mechanisms emerge in architectures incorporating recurrence, attention, or feedback. Second, our training regime implemented an extreme imbalance (100% Asian vs. 100% White faces), future work should examine more graded imbalances (e.g., 80/20 distributions) and variations in exposure quality (e.g., many identities collapsed into a single category) to further differentiate theoretical accounts of the ORE (58). Finally, when relating models to human behavior, we focused on observers whose everyday experience is skewed toward their own facial group and did not include individuals with substantial, balanced exposure to both groups. Consequently, we could not directly test whether the Dual CNN best captures behavior in observers with genuinely diverse face experience.

In summary, our study shows that training diversity in CNNs enhances recognition performance across groups and promotes integrated, generalizable representational geometries that more closely resemble human perceptual organization. By combining behavioral alignment with representational similarity analysis and lesioning, we show that diverse experience mitigates group-specific overfitting, improves identity discrimination, and yields an integrated feature space rather than segregated group-specific subsystems, allowing faces from different groups to be encoded within a shared representational geometry. More broadly, our results underscore the central role of exposure diversity in shaping both artificial and biological face-processing systems and highlight representational diversity as essential for building accurate, equitable, and cognitively plausible AI models. Together, these contributions move beyond documenting ORE in CNNs and toward a computational account of its origins, its representational consequences, and the conditions under which it can be reduced.

## Methods

### Training Deep Convolutional Neural Networks

To assess the impact of training diversity on face representations and recognition performance, we trained three CNNs, all based on the VGG16 architecture (40; Figure 4A). VGG16 was chosen because it is optimized for face recognition (57), achieves human-level accuracy on diverse face recognition tasks (59, 60) and reproduces behavioral and neural signatures of human face perception (29, 61).

Each network was randomly initialized and trained for identity recognition using datasets that differed in facial composition (Figure 4B): First, a Single Asian CNN was trained exclusively on 1,654 Asian identities (891 female) from a subset of the Asian Face Dataset (62). Second, we trained a Single White CNN exclusively on 1,654 White identities (827 female) drawn from subsets of the VGGFace2 (63) and CASIA-WebFace (64) datasets. For both the single-trained CNNs, 94 training images and 5 validation images per identity were used. The number of identities and images per identity was matched across groups and was constrained by the available size of the Asian dataset, ensuring that differences between models reflect training diversity rather than differences in dataset size. Third, we trained a Dual CNN on the union of both groups (3,308 identities total). To equate the total number of training samples across all models (155,476 total training images), we used 47 training images per identity (validation images remained 5 per identity).

All three CNNs were trained using identical hyperparameters. Training was conducted with a batch size of 128, using a cross-entropy loss function and the stochastic gradient descent (SGD) optimizer with a momentum of 0.9 and a learning rate of 0.001. To prevent overfitting, a weight decay of 0.0001 was utilized. A learning rate scheduler (ReduceLROnPlateau, PyTorch) reduced the learning rate by a factor of 0.1 if the validation loss failed to decrease for five consecutive epochs (‘patience’ = 5). This helped fine-tune learning when training reached a plateau.

During training, all images underwent a series of data augmentations. Each image was resized to 256 x 256 pixels and then randomly cropped to 224 x 224 pixels. A random grayscale transformation was applied with a probability of 0.2, ensuring that the networks learned features beyond color cues. Additionally, 50% of the images were randomly flipped horizontally to increase variability. Finally, all images were normalized to a mean of 0.5 and a standard deviation of 0.5 for each RGB channel. For validation and testing, images were resized to 256 x 256 pixels, center-cropped to 224 x 224 pixels, and normalized using the same parameters.

### Other-race Effect in CNNs

To assess whether the ORE was present in the CNNs, we evaluated each CNN on a target-matching task using novel faces not included during training (Figure 4C). The test set consisted of young (20-35 years), female White and Asian identities. Specifically, we used a dataset of 40 Asian and 40 White identities, each with 5 images (200 images per group; 400 images total) (29). The images were collected from colleagues or public Instagram accounts (< 2,000 followers). To control for low-level differences between face groups, we quantified grayscale pixel variance for each image and compared the averages across groups. Images were first converted to greyscale, and intensity values were scaled to the range [0,1]. Pixel variance was then computed across all pixels within each image, yielding one variance value per image. Pixel variance was compared between facial groups using a permutation test. For each group, we generated 1,560 triplets (40 × 39), consisting of two images of a target identity and one distractor image from a different identity. Performance was quantified based on the cosine similarity between image feature vectors from the penultimate fully connected layer (fc7) of each CNN. We chose this layer because it captures the model’s learned identity-relevant features without being tied to the specific output units (65). For each triplet, the image with the higher cosine similarity to the target was taken as the model’s choice (Figure 4C).

**ORE Index**: To quantify recognition bias across facial groups for all CNNs, we defined a difficulty-weighted ORE index based on normalized accuracies. First, we estimated a group-specific performance ceiling using the trained group accuracy of the corresponding single-trained model:

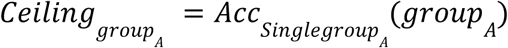

For each model *m*, we then computed normalized accuracies:

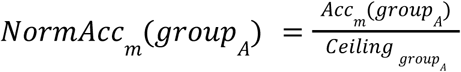

Finally, we defined the difficulty-weighted ORE index as:

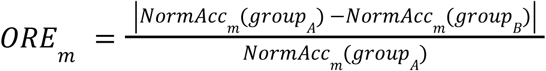

For the Dual CNN, the ORE index was computed in both directions, allowing us to assess the symmetry of bias across groups. This normalization procedure corrects for inherent differences in group-specific task difficulty, ensuring that any observed ORE differences across models reflect true representational bias rather than differences in baseline recognizability arising from the distinct stimulus sets.

### Obtaining Properties of the Representational Feature Space

To investigate the properties of the learned representational feature space across CNNs, we used three complementary approaches. First, we applied a lesioning analysis in the Dual CNN to test whether diverse training experience led to an integrated feature space or to group-specific subspaces within the network. Second, we performed Representational Similarity Analysis (RSA; 66) to compare the internal representational geometries across all CNNs. Finally, we visualized filter-preferred stimuli by iteratively optimizing random noise images to maximize the activation of filters most selective for faces. This provided qualitative insight into how training affected feature preferences in each CNN.

#### Lesioning Analysis in the Dual CNN

To test whether the processing of different facial groups (Asian and White faces) was segregated or integrated within the last convolutional layer (conv13) of the Dual CNN, we applied a lesioning analysis following the procedure described in Dobs et al. (2023). We focused on conv13 because it is the final layer containing spatially localized filters, thus making it the last point at which feature selectivity can be meaningfully attributed to individual filters. We first obtained group-specific importance rankings by ablating each filter in the conv13 individually and measuring the resulting increase in loss separately for each group, using 50 batches of training images per group. Filters were then ranked independently for the Asian and White face-recognition tasks based on their respective contributions to the loss.

From this initial ranking, we performed progressive, conditional lesioning in 1.6% increments: 1) Lesion the top 1.6% of filters (for a given group) simultaneously and keep them lesioned. 2) Recompute the individual loss ranking on the remaining filters (with previously selected filters still lesioned). 3) Add the top 1.6% from this updated ranking to the lesioned set and keep them lesioned in the next iteration. We repeated steps (2)-(3) until 20% of filters were selected for each group. This procedure conditions each selection on prior lesions, reducing redundancy and capturing filters that remain most informative given previously removed filters.

Finally, for each group-specific set, we lesioned the full 20% of filters and evaluated performance on the entire validation set for both groups, allowing us to test whether the Dual CNN relies on group-specific subsets (segregated subspaces) or a shared, integrated feature space. We chose a 20% lesion proportion because prior work (33) found that this level of ablation produces strong and interpretable task-specific effects while avoiding excessive overlap between critical filters across tasks.

#### Representational Similarity Analysis

To characterize the representational geometry of each CNN, we conducted Representational Similarity Analysis (RSA) using the same stimulus set of 400 images (40 identities × 5 images each; Asian and White faces) used to evaluate the ORE. Activations from the fc7 layer were extracted for all images, and pairwise dissimilarities were computed using 1 – cosine similarity, resulting in a 400 x 400 representational dissimilarity matrix (RDM) for each model.

To evaluate how training diversity shapes representational geometries, we quantified dissimilarities within facial groups (Asian–Asian, White–White) and between groups (Asian–White) for each CNN. We additionally visualized the RDMs using Multidimensional Scaling (MDS) to obtain lower-dimensional embeddings that qualitatively reveal clustering and separation patterns.

Finally, we performed cross-model RSA by correlating RDMs across networks using the Pearson correlation coefficient to assess representational similarity between CNNs trained on different datasets. Together, these analyses allowed us to directly compare how differently trained CNNs encode facial identity and group structure in their representational spaces.

### Comparing Human Participants and CNNs

To directly assess the correspondence between CNN performance and human behavior, we compared the decision patterns of the CNNs with those of Asian (*n* = 102) and White (*n* = 269) participants from a previously published study (29). Human responses were obtained using the same target-matching task illustrated in Figure 4D, in which participants selected which of two faces matched the target identity. This allowed a direct comparison between CNNs and human decision patterns across facial groups and stimulus conditions. The ORE was quantified as the proportional reduction in recognition accuracy for other-race relative to own-race faces, with larger values indicating a stronger effect. For details on participant recruitment, experimental procedures, and data collection, see Dobs et al. (2023).

#### Stimuli and Target-matching Task in Humans

The stimuli used for comparing the human and CNNs choices were identical to those used for evaluating the ORE and behavioral data were drawn from Dobs et al. (2023). Each participant completed 40 target-matching trials (20 per facial group) presented in randomized order, in which they chose which of two faces matched a target identity (Figure 4D). For detailed information on the experimental setup, please refer to Dobs et al. (2023).

To assess the reliability of participants’ choices, we conducted a split-half reliability analysis across triplets. In each of 1,000 iterations, we randomly divided participants into two independent groups. For each group, we computed average triplet-choice patterns and retained those triplets that were shared between both groups to ensure valid comparisons. We then calculated Pearson correlation coefficients between the two halves and applied the Spearman-Brown prophecy correction to correct for biases introduced by split-half calculations. We used the formula 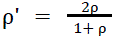 where ⍴ is the Pearson correlation between the two halves. To express the corrected reliability on the same scale as Pearson correlation coefficients, we report the square root of ⍴’. Note that because each participant viewed only a small subset of all triplets, not all triplets were shared across participants, which likely led to a slight underestimation of the true noise ceiling.

#### CNN-Human Similarity Analysis

We then quantified the correspondence between CNN and human decision patterns at the triplet level. For each CNN, activations for the triplets (Target, Match A and Match B) used in the human experiment were extracted from the fc7 layer. We selected fc7 because it is the penultimate layer, proximal to the network’s decision output yet not directly constrained by the classification layer, thus providing a meaningful approximation of the model’s representational behavior. Cosine similarities between each match and the target were calculated and the probability of choosing Match A was obtained from:

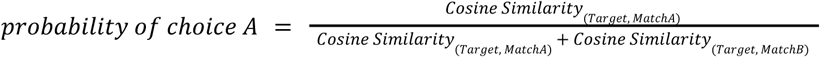

Next, the probabilities of correct choices for triplets were computed as averages across participants. Finally, these behavioral choice probabilities were correlated with CNNs’ choice probabilities obtained from the previous step using a Pearson correlation. This correlation quantifies the level of correspondence between model and human decision patterns.

### Statistical Inference

Nonparametric statistical methods were employed to avoid assumptions about data distribution. For the ORE analysis, we bootstrapped the 1,560 triplets 10,000 times. For each resample, average identity-matching accuracy was computed, resulting in a distribution of ORE values for each CNN. We defined 95% confidence intervals as the 2.5th and 97.5th percentiles of the bootstrapped distributions. Pairwise comparisons between CNNs were performed separately for each facial group (Asian and White faces), with two-tailed p-values derived from the bootstrapped distribution of accuracy differences. All p-values were adjusted for multiple comparisons (two pairwise contrasts) using the Bonferroni correction, with a significance threshold of *p* < .05.

For the CNN-human consistency analysis, we calculated Pearson correlation coefficients between CNN and human responses across triplets within each bootstrap sample. Distributions of correlation values were compared between CNNs, stratified by both stimulus group (Asian vs. White faces) and participant group (Asian vs. White participants). Error bars indicate the variability of correlation coefficients, quantified as the standard deviation across bootstrap samples. To compare the consistency of two CNNs, as in the ORE analysis, two-tailed p-values were obtained from bootstrapped differences. All p-values were adjusted for multiple comparisons (three pairwise contrasts) using the Bonferroni correction, with a significance threshold of *p* < .05.

## Data and Code Availability

All analyses were conducted using Python 3.9.12, and model training was performed using Python 3.8 and PyTorch 1.13.0. All analysis scripts are available at https://github.com/VCCN-lab/ORE_in_CNNs_and_humans. Code for training the networks is available at https://github.com/martinezjulio/sdnn (33). The trained CNN models will be made available upon publication of the paper. All stimuli used for computing ORE, RSA, and for human-model behavioral comparisons are available at OSF (https://osf.io/dbks3/) (29).

## Acknowledgements

We thank the members of the VCCN lab for valuable feedback and discussion, and Ben de Haas for comments on an earlier version of the manuscript. This work was supported by the European Research Council (ERC Starting Grant DEEPFUNC, ERC-2023-STG-101117441), the Hessian Ministry of Higher Education, Research, Science and the Arts (LOEWE Start Professorship), and the Deutsche Forschungsgemeinschaft (DFG, German Research Foundation) under Germany’s Excellence Strategy (EXC 3066/1 “The Adaptive Mind”, Project No. 533717223) and SFB/TRR 135 (Project No. 222641018, TP C9).

## References

1. C. A. Meissner, J. C. Brigham, Thirty years of investigating the own-race bias in memory for faces: A meta-analytic review. Psychol. Public Policy Law 7, 3–35 (2001).

2. S. J. Platz, H. M. Hosch, Cross-Racial/Ethnic Eyewitness Identification: A Field Study^1^. J. Appl. Soc. Psychol. 18, 972–984 (1988).

3. B. Rossion, C. Michel, An experience-based holistic account of the other-race face effect. The Oxford handbook of face perception (Oxford University Press, 2012).

4. D. J. Turk, T. C. Handy, M. S. Gazzaniga, Can perceptual expertise account for the own-race bias in face recognition? A split-brain study. Cogn. Neuropsychol. 22, 877–883 (2005).

5. P. M. Walker, M. Hewstone, A perceptual discrimination investigation of the own-race effect and intergroup experience. Appl. Cogn. Psychol. 20, 461–475 (2006).

6. P. M. Walker, J. W. Tanaka, An Encoding Advantage for Own-Race versus Other-Race Faces. Perception 32, 1117–1125 (2003).

7. H. K. Wong, A. J. Estudillo, I. D. Stephen, D. R. T. Keeble, The other-race effect and holistic processing across racial groups. Sci. Rep. 11, 8507 (2021).

8. M. Zhao, W. G. Hayward, I. Bulthoff, Face format at encoding affects the other-race effect in face memory. J. Vis. 14, 6–6 (2014).

9. L. Ficco, V. I. Müller, J. M. Kaufmann, S. R. Schweinberger, Socio-cognitive, expertise-based and appearance-based accounts of the other-‘race’ effect in face perception: A label-based systematic review of neuroimaging results. Br. J. Psychol. 114, 45–69 (2023).

10. A. J. Golby, J. D. E. Gabrieli, J. Y. Chiao, J. L. Eberhardt, Differential responses in the fusiform region to same-race and other-race faces. Nat. Neurosci. 4, 845–850 (2001).

11. D. J. Kelly, et al., Three-month-olds, but not newborns, prefer own-race faces. Dev. Sci. 8 (2005).

12. Y. Bar-Haim, T. Ziv, D. Lamy, R. M. Hodes, Nature and Nurture in Own-Race Face Processing. Psychol. Sci. 17, 159–163 (2006).

13. D. J. Kelly, et al., Cross-Race Preferences for Same-Race Faces Extend Beyond the African Versus Caucasian Contrast in 3-Month-Old Infants. Infancy 11, 87–95 (2007).

14. D. J. Kelly, et al., The Other-Race Effect Develops During Infancy: Evidence of Perceptual Narrowing. Psychol. Sci. 18, 1084–1089 (2007).

15. G. Anzures, et al., Brief daily exposures to Asian females reverses perceptual narrowing for Asian faces in Caucasian infants. J. Exp. Child Psychol. 112, 484–495 (2012).

16. G. Anzures, et al., Developmental Origins of the Other-Race Effect. Curr. Dir. Psychol. Sci. 22, 173–178 (2013).

17. E. McKone, et al., A critical period for faces: Other-race face recognition is improved by childhood but not adult social contact. Sci. Rep. 9, 12820 (2019).

18. S. M. Spangler, et al., The Other-Race Effect in a Longitudinal Sample of 3-, 6– and 9-Month-Old Infants: Evidence of a Training Effect. Infancy 18, 516–533 (2013).

19. J. Suhrke, et al., The other-race effect in 3-year-old German and Cameroonian children. Front. Psychol. 5 (2014).

20. L. Wan, K. Crookes, K. J. Reynolds, J. L. Irons, E. McKone, A cultural setting where the other-race effect on face recognition has no social–motivational component and derives entirely from lifetime perceptual experience. Cognition 144, 91–115 (2015).

21. J. C. Brigham, R. S. Malpass, The Role of Experience and Contact in the Recognition of Faces Of Own– and Other-Race Persons. J. Soc. Issues 41, 139–155 (1985).

22. K. J. Hancock, G. Rhodes, Contact, configural coding and the other-race effect in face recognition. Br. J. Psychol. 99, 45–56 (2008).

23. A. J. O’Toole, K. Deffenbacher, H. Abdi, J. C. Bartlett, Simulating the ‘Other-race Effect’ as a Problem in Perceptual Learning. Connect. Sci. 3, 163–178 (1991).

24. P. M. Walker, L. Silvert, M. Hewstone, A. C. Nobre, Social contact and other-race face processing in the human brain. Soc. Cogn. Affect. Neurosci. 3, 16–25 (2008).

25. B. Balas, A. Saville, Hometown size affects the processing of naturalistic face variability. Vision Res. 141, 228–236 (2017).

26. G. Jacob, R. T. Pramod, H. Katti, S. P. Arun, Qualitative similarities and differences in visual object representations between brains and deep networks. Nat. Commun. 12, 1872 (2021).

27. N. Kriegeskorte, Deep Neural Networks: A New Framework for Modeling Biological Vision and Brain Information Processing. Annu. Rev. Vis. Sci. 1, 417–446 (2015).

28. D. L. K. Yamins, J. J. DiCarlo, Using goal-driven deep learning models to understand sensory cortex. Nat. Neurosci. 19, 356–365 (2016).

29. K. Dobs, J. Yuan, J. Martinez, N. Kanwisher, Behavioral signatures of face perception emerge in deep neural networks optimized for face recognition. Proc. Natl. Acad. Sci. 120, e2220642120 (2023).

30. G. Yovel, I. Grosbard, N. Abudarham, Deep learning models of perceptual expertise support a domain-specific account. [Preprint] (2022). http://biorxiv.org/lookup/doi/10.1101/2022.12.01.518342 (Accessed 15 September 2025).

31. A. Zeman, T. Leers, H. O. De Beeck, Mooney Face Image Processing in Deep Convolutional Neural Networks Compared to Humans. [Preprint] (2022). http://biorxiv.org/lookup/doi/10.1101/2022.03.21.485240 (Accessed 15 September 2025).

32. J. Tian, H. Xie, S. Hu, J. Liu, Multidimensional Face Representation in a Deep Convolutional Neural Network Reveals the Mechanism Underlying AI Racism. Front. Comput. Neurosci. 15, 620281 (2021).

33. K. Dobs, J. Martinez, A. J. E. Kell, N. Kanwisher, Brain-like functional specialization emerges spontaneously in deep neural networks. Sci. Adv. 8, eabl8913 (2022).

34. N. Abudarham, I. Grosbard, G. Yovel, Face Recognition Depends on Specialized Mechanisms Tuned to View-Invariant Facial Features: Insights from Deep Neural Networks Optimized for Face or Object Recognition. Cogn. Sci. 45 (2021).

35. M. Rosemblaum, et al., Concurrent emergence of view invariance, sensitivity to critical features, and identity face classification through visual experience: Insights from deep learning algorithms. J. Vis. 25, 2 (2025).

36. A. Doerig, et al., The neuroconnectionist research programme. Nat. Rev. Neurosci. 24, 431–450 (2023).

37. N. Kanwisher, M. Khosla, K. Dobs, Using artificial neural networks to ask ‘why’ questions of minds and brains. Trends Neurosci. 46, 240–254 (2023).

38. A. J. O’Toole, C. D. Castillo, Face Recognition by Humans and Machines: Three Fundamental Advances from Deep Learning. Annu. Rev. Vis. Sci. 7, 543–570 (2021).

39. L. E. Van Dyck, W. R. Gruber, Modeling Biological Face Recognition with Deep Convolutional Neural Networks. J. Cogn. Neurosci. 35, 1521–1537 (2023).

40. K. Simonyan, A. Zisserman, Very Deep Convolutional Networks for Large-Scale Image Recognition. [Preprint] (2015). http://arxiv.org/abs/1409.1556 (Accessed 7 April 2025).

41. A. Wang, M. W. Sliwinska, D. M. Watson, S. Smith, T. J. Andrews, Distinct patterns of neural response to faces from different races in humans and deep networks. Soc. Cogn. Affect. Neurosci. 18, nsad059 (2023).

42. N. Kanwisher, G. Yovel, The fusiform face area: a cortical region specialized for the perception of faces. Philos. Trans. R. Soc. B Biol. Sci. 361, 2109–2128 (2006).

43. N. Kanwisher, J. McDermott, M. M. Chun, The Fusiform Face Area: A Module in Human Extrastriate Cortex Specialized for Face Perception. J. Neurosci. 17, 4302–4311 (1997).

44. T. Valentine, A Unified Account of the Effects of Distinctiveness, Inversion, and Race in Face Recognition. Q. J. Exp. Psychol. Sect. A 43, 161–204 (1991).

45. M. Shoura, Y. Z. Liang, M. A. Sama, A. De, A. Nestor, Revealing the neural representations underlying other-race face perception. Front. Hum. Neurosci. 19, 1543840 (2025).

46. L. Vizioli, G. A. Rousselet, R. Caldara, Neural repetition suppression to identity is abolished by other-race faces. Proc. Natl. Acad. Sci. 107, 20081–20086 (2010).

47. U. Cohen, S. Chung, D. D. Lee, H. Sompolinsky, Separability and geometry of object manifolds in deep neural networks. Nat. Commun. 11, 746 (2020).

48. J. Mehrer, C. J. Spoerer, E. C. Jones, N. Kriegeskorte, T. C. Kietzmann, An ecologically motivated image dataset for deep learning yields better models of human vision. (2021).

49. M. Lehtonen, V. Fyndanis, J. Jylkkä, The relationship between bilingual language use and executive functions. Nat. Rev. Psychol. 2, 360–373 (2023).

50. H. K. Wong, I. D. Stephen, D. R. T. Keeble, The Own-Race Bias for Face Recognition in a Multiracial Society. Front. Psychol. 11, 208 (2020).

51. J. W. Tanaka, B. Heptonstall, S. Hagen, Perceptual expertise and the plasticity of other-race face recognition. Vis. Cogn. 21, 1183–1201 (2013).

52. M. J. Bernstein, S. G. Young, K. Hugenberg, The Cross-Category Effect: Mere Social Categorization Is Sufficient to Elicit an Own-Group Bias in Face Recognition. Psychol. Sci. 18, 706–712 (2007).

53. E. R. Shriver, S. G. Young, K. Hugenberg, M. J. Bernstein, J. R. Lanter, Class, Race, and the Face: Social Context Modulates the Cross-Race Effect in Face Recognition. Pers. Soc. Psychol. Bull. 34, 260–274 (2008).

54. S. G. Young, K. Hugenberg, M. J. Bernstein, D. F. Sacco, Perception and Motivation in Face Recognition: A Critical Review of Theories of the Cross-Race Effect. Personal. Soc. Psychol. Rev. 16, 116–142 (2012).

55. I. D. Raji, G. Fried, About Face: A Survey of Facial Recognition Evaluation. (2021).

56. R. Singh, P. Majumdar, S. Mittal, M. Vatsa, Anatomizing Bias in Facial Analysis. [Preprint] (2021). http://arxiv.org/abs/2112.06522 (Accessed 16 April 2025).

57. O. M. Parkhi, A. Vedaldi, A. Zisserman, Deep Face Recognition in Procedings of the British Machine Vision Conference 2015, (British Machine Vision Association, 2015), p. 41.1-41.12.

58. M. K. Qian, et al., Perceptual individuation training (but not mere exposure) reduces implicit racial bias in preschool children. Dev. Psychol. 53, 845–859 (2017).

59. T. Gwyn, K. Roy, M. Atay, Face Recognition Using Popular Deep Net Architectures: A Brief Comparative Study. Future Internet 13, 164 (2021).

60. M. Wang, W. Deng, Deep face recognition: A survey. Neurocomputing 429, 215–244 (2021).

61. S. Grossman, et al., Convergent evolution of face spaces across human face-selective neuronal groups and deep convolutional networks. Nat. Commun. 10, 4934 (2019).

62. Z. Xiong, et al., “An Asian Face Dataset and How Race Influences Face Recognition” in Advances in Multimedia Information Processing – PCM 2018, Lecture Notes in Computer Science., R. Hong, W.-H. Cheng, T. Yamasaki, M. Wang, C.-W. Ngo, Eds. (Springer International Publishing, 2018), pp. 372–383.

63. Q. Cao, L. Shen, W. Xie, O. M. Parkhi, A. Zisserman, VGGFace2: A dataset for recognising faces across pose and age. [Preprint] (2018). http://arxiv.org/abs/1710.08092 (Accessed 7 April 2025).

64. D. Yi, Z. Lei, S. Liao, S. Z. Li, Learning Face Representation from Scratch. [Preprint] (2014). http://arxiv.org/abs/1411.7923 (Accessed 7 April 2025).

65. A. J. O’Toole, C. D. Castillo, C. J. Parde, M. Q. Hill, R. Chellappa, Face Space Representations in Deep Convolutional Neural Networks. Trends Cogn. Sci. 22, 794–809 (2018).

66. N. Kriegeskorte, M. Mur, P. Bandettini, Representational similarity analysis – connecting the branches of systems neuroscience. Front. Syst. Neurosci. (2008). 10.3389/neuro.06.004.2008.

